# FMRP regulates adult human cortical neuron excitability via cyclic-AMP signalling

**DOI:** 10.1101/2025.10.14.682273

**Authors:** Max JJ Knops, Soraya Meftah, Max A Wilson, Lewis W Taylor, Calum Bonthron, Alsadeg Bilal, Imran Liaquat, Paul M Brennan, Claire S Durrant, Sam A Booker

**Affiliations:** Institute for Neuroscience and Cardiovascular Research (INCR), University of Edinburgh, Edinburgh, UK; Simons Initiative for the Developing Brain, University of Edinburgh, Edinburgh, UK; Patrick Wild Centre for Research into Fragile X Syndrome, Autism, and intellectual disability, University of Edinburgh, Edinburgh, UK; UK Dementia Research Institute, University of Edinburgh, Edinburgh, UK; Department of Clinical Neurology, NHS Lothian, Royal Infirmary Edinburgh, UK

**Keywords:** Fragile X Syndrome, *FMR1*, FMRP, hyperexcitability, human, slice culture, phosphodiesterase 4D, BPN-14, 770

## Abstract

Fragile X Syndrome (FXS) is a common inherited neurodevelopmental condition, resulting from loss of Fragile X Messenger Ribonuclear Protein (FMRP). Rodent models of FXS display cellular hyperexcitability, but it is not known to what extent this is the case in intact human neurons. Depleting FMRP in human brain slice cultures reveals cyclic-AMP-dependent cellular hyperexcitability which is corrected by phosphodiesterase 4D inhibition and may be independent of neurodevelopment.

## Main text

Fragile X Syndrome (FXS) has been identified as a condition of elevated neuronal excitability, with patients displaying enhanced cortical EEG signals ^1^, and with cellular correlates of such observed in rodent ^2-4^ and induced pluripotent stem-cell (iPSC) models ^5^; which occur alongside changes in protein synthesis ^6^. Such excitability has been linked to impaired neuronal homeostasis ^7^, direct interactions of ion channels with Fragile X Messenger Ribonuclear Protein (FMRP) and altered protein phosphorylation ^8^. Alongside this range of functions, loss of FMRP has been implicated with reduced cyclic adenosine monophosphate (cAMP) levels in humans, rodent and invertebrate models, and iPSC lines^9^. This is exemplified by phosphodiesterase (PDE) inhibitors being identified as a promising target for the treatment of FXS, particularly the brain-specific 4D isoform ^10^. However, PDE4D inhibitors display human specificity ^11^, limiting their functional validation in other models. These models also do not fully recapitulate the complexity of the postnatal human brain ^12^. It remains unknown how FMRP regulates neuron function in intact human brain tissue, and whether loss of FMRP is amenable to therapies targeting PDE4D in humans.

We hypothesised that loss of FMRP would lead to cellular hyperexcitability in adult neurons, which may be amenable to PDE4D inhibition. To address this, we have developed a living human brain tissue model of FXS, in which we cultured adult human brain slices for up to 12 days *in vitro* (DIV)^13^. For the present study, we report data from brain tissue from 23 patients collected during surgical removal of brain tumours (48% glioblastoma, 43% glioma, 9% other), none of whom had been diagnosed with FXS. The median age of patients was 61 years (range: 23 – 79) comprising 83% biological male and 17% biological female individuals. 61% of patients had not experienced seizures, meanwhile 39% had experienced a seizure and been prescribed levetiracetam (1g/day; Supplementary Table 1).

To deplete FMRP, we applied a lentiviral vector expressing a short-hairpin RNA recognising human *FMR1* (*FMR1*-shRNA) or *Scrambled* controls, both with a green-fluorescent protein (GFP) reporter (Figure 1A). We observe consistent GFP expression across the cortex of cultured slices at 8-10 DIV (Figure 1B). Analysis of immunolabelling revealed that FMRP levels are substantially reduced in *FMR1*-shRNA cultures (Figure 1C), compared with non-transduced GFP-negative neurons or *Scrambled* controls (Figure 1D), with an average 32% reduction in FMRP immunofluorescence (Figure 1E). The U6 promoter we used targets all cells. Overall, we found that in MAP2-immunoreactive neurons, 48.2 ± 9.6% of cells in *Scrambled* and 38.8 ± 9.9 % in *FMR1*-shRNA treated cultures were GFP positive (Figure 1F). Conversely, of GFP-labelled transduced cells, 34.4 ± 6.7% of *Scrambled* and 25.6 ± 5.3 % of *FMR1*-shRNA treated cells were MAP2 positive (Figure 1G). These data confirm that we have established a human slice culture model in which we reliably deplete FMRP; which displays a mosaic pattern in neurons.

**Figure 1.**
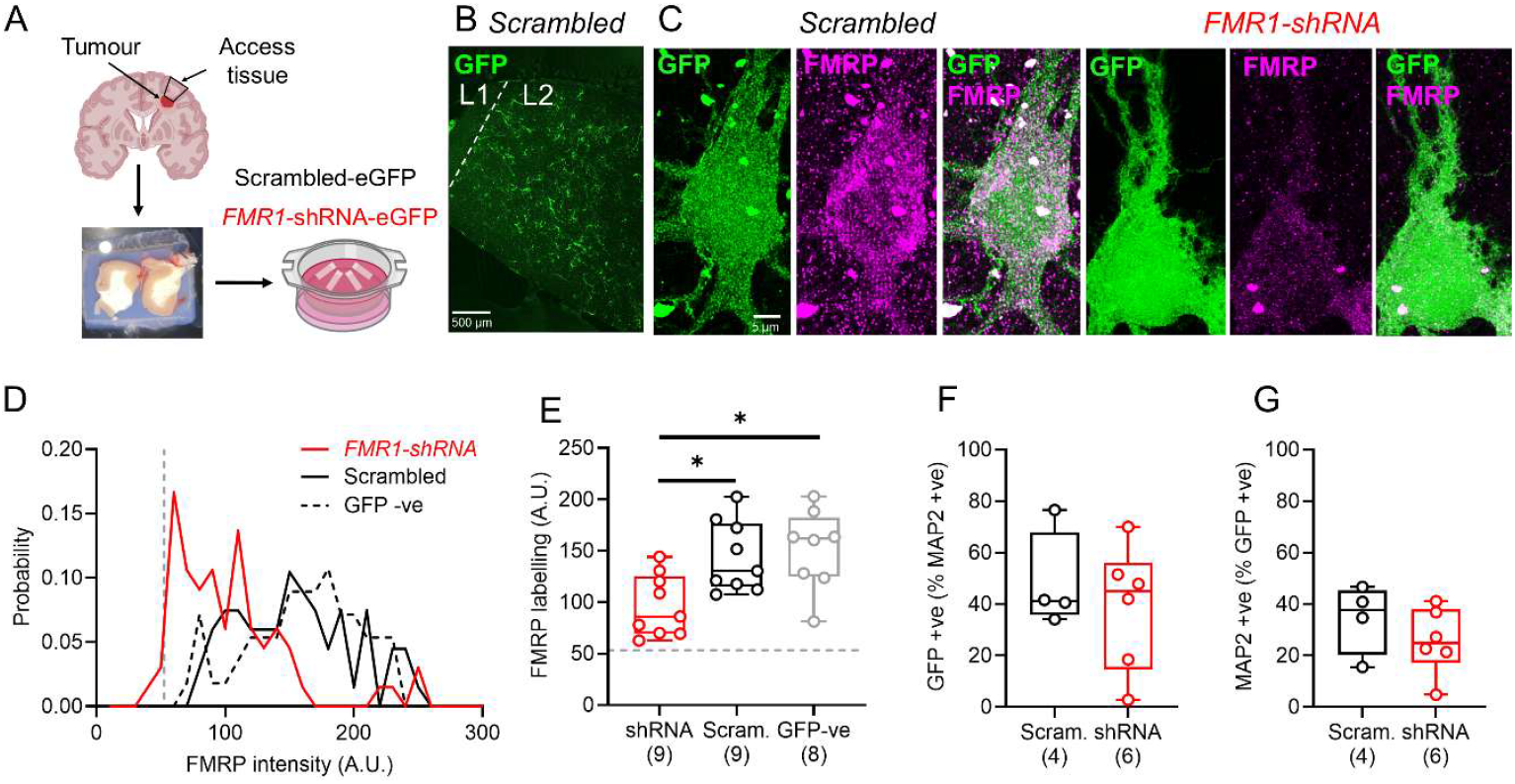
Transduction of adult human slice cultures with FMR1-shRNA leads to depletion of FMRP in neurons. A) Overview of experimental pipeline. B) Example low-magnification flattened confocal stack of a human slice culture transduced with *Scrambled* GFP-expressing lentivirus (green), in L1 and L2/3 of neocortex (dashed line). C) Example images of FMRP immunostaining (magenta) in GFP expressing (green) cells in human slice cultures transduced with either *Scrambled* (left) or *FMR1-* shRNA (right), including merge of both signals. D) Probability distribution of FMRP fluorescence intensity (arbitrary units [A.U.]) measured at the soma of cells transduced with *FMR1*-shRNA (red lines) or *Scrambled* (black lines) lentiviral constructs, compared to GFP-negative cells (black dashed lines) and background fluorescence (grey dashed line). E) Relative FMRP fluorescence of tested cultures (open circles) measuring *FMR1*-shRNA (red), *Scrambled* (black) or GFP-negative cells (grey). The grey dashed line indicates the average background intensity. F) percentage of MAP2-immunoreactive cells that also expressed GFP in *FMR1*-shRNA and *Scrambled* cultures. G) Percentage of GFP-positive cells that expressed MAP2. Data is shown as either probability distributions (D) or box plots, depicting the median with 25-75% quartile range, and maximum & minimum values. Data from individual cultures are shown overlaid (E, F, G) and number of patients indicated in parentheses. All statistical tests were performed using a LMM (Var∼Genotype +(1|Brain Region)+(1|Case/Slice). with type-3 ANOVA, shown as: * – *p*<0.05. Some elements created with BioRender.

Loss of FMRP in rodent and iPSC models leads to elevated neuronal excitability ^8,14^, in particular impaired ion channel function contributes to membrane excitability and action potential (AP) discharge ^2,15-17^. To determine whether loss of FMRP in adult human slice culture neurons impairs neuronal excitability we next performed whole-cell patch-clamp recordings from GFP-positive neurons from L2/3 of cultured slices at DIV7-14 transduced with either *FMR1*-shRNA or *Scrambled* constructs. In both cultures we observed cells that produced APs (AP) in response to depolarising stimuli (Figure 2A). As our viral vector utilised the ubiquitous U6 promoter we only assessed cells that displayed APs (neurons), had a membrane potential more hyperpolarised than -40 mV and an input resistance less than 1000 mΩ ^18^. We observed that neurons expressing *Scrambled* constructs tended to fire few APs, consistent with previous reports ^19^. *FMR1-*shRNA neurons, by contrast, displayed higher AP discharge for the same current steps (Figure 2B). We found minimal change in passive properties such as resting membrane potential, input resistance, impedance, or resonant frequency (Figure 2C). The enhanced neuronal firing appeared to be due to a number of changes in neuronal physiology, notably reduced rheobase, hyperpolarised voltage threshold, alongside accelerated AP maximum rise and decay rates (Figure 2D). A full summary of electrophysiological parameters measures is presented in Supplementary Table 2. Similar patterns of activity were observed in all recorded cells, regardless of resting membrane potential (Supplementary Figure 1). We found no effect of biological sex or seizures history, but while sampled brain region did have an effect on neuronal excitability, with frontal and temporal cortices displaying the highest firing rates; enhanced firing was consistently observed following loss of FMRP (Supplementary Figure 2).

**Figure 2.**
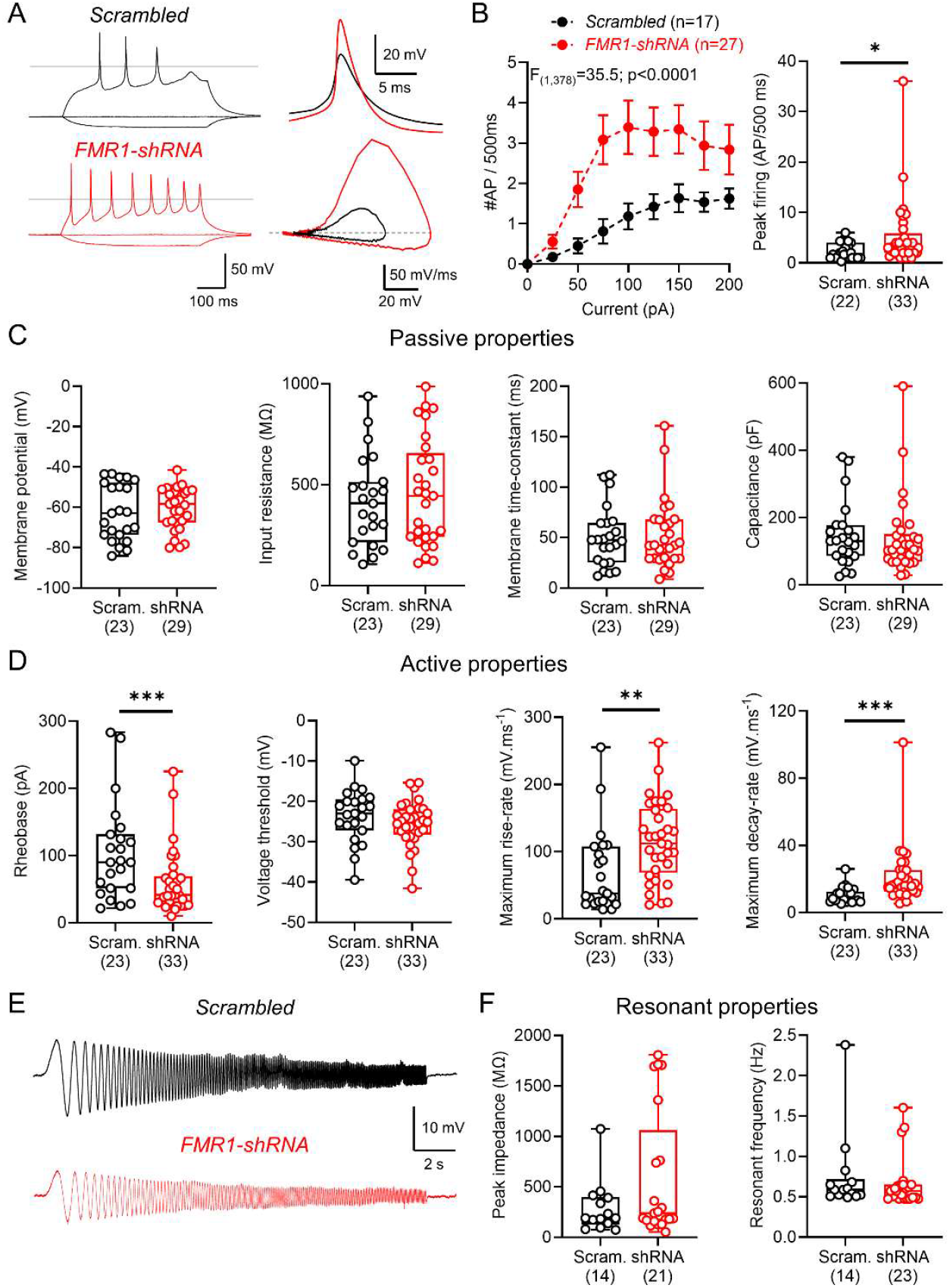
Loss of FMRP in adult human slice culture neurons increases neuronal excitability due to enhanced voltage-activated currents. A) Left, example voltage responses of neurons in human slice cultures from *Scrambled* (black) or *FMR1*-shRNA (red) transduced neurons in response to -25, 0, and + 100 pA current steps (500 ms). Right, example AP waveforms from *Scrambled* and *FMR1*-shRNA transduced neurons and the phase plots of the same responses. B) AP output to depolarising current steps from human slice culture neurons recorded with a resting membrane potential below -40 mV. C) Quantification of passive membrane properties of recorded neurons, including resting membrane potential, input resistance, membrane time-constant, and capacitance. D) Measurement of active properties of neurons, including rheobase current, voltage threshold, maximum AP rise-rate, and maximum AP decay rate. E) Example voltage responses to a 50 pA (peak-to-trough) frequency-modulated (0.1 – 20 Hz, 20 s) sinusoidal wave from *Scrambled* (black) or *FMR1*-shRNA (red) transduced neurons. F) Peak impedance and resonant frequency of measured from sinusoidal stimuli in recorded neurons. Data is shown as either mean ± SEM (B) or box plots, depicting the median with 25-75% quartile range, and maximum & minimum. Data from individual transduced neurons shown overlaid and the number of cells shown in parenthesis, from 8 patients (*Scrambled*) or 12 patients (*FMR1*-shRNA). All statistical tests were performed using 2-way ANOVA (B) and LMM or GLMM with type-3 ANOVA and shown as: n.s – *p*>0.05, * – *p*<0.05, ** – *p*>0.01, *** – *p*<0.001, or reported above graphs.

As many studies have indicated a synaptic basis of FXS (reviewed in Contractor, et al. ^20^), we also compared the synaptic inputs to *Scrambled* and *FMR1*-shRNA neurons. Spontaneous excitatory postsynaptic currents did not differ in their amplitude or frequency (Supplementary Figure 3), consistent with findings in rodent models of FXS from neocortex ^2^ and hippocampus ^21^. We found that depletion of FMRP for 8-14 DIV did lead to truncated dendritic arbours in recorded neurons, when measured in a subset of recorded neurons (Supplementary Figure 3D-F).

Overall, these data reveal that depletion of FMRP from adult human neurons *in vitro* leads to increased neuronal excitability, which appears to be due to changes in AP kinetics. These data indicate a lifelong role for FMRP to maintain the excitability of human neurons, independent of the neurodevelopmental program.

A key function of FMRP is regulating cAMP levels, as plasma levels are reduced in affected individuals, and iPSC and rodent models of FXS. Increasing cAMP, therefore, may reflect a plausible treatment option for FXS^9^. Indeed, the PDE4D inhibitor BPN-14,770, boosts cAMP levels both *in vitro* and *in vivo*, with greater affinity for human isoforms leading to enhanced circuit function^11^ and has shown promise in clinical trials for FXS^22^. To determine whether PDE4D inhibition is capable of correcting cellular hyperexcitability in *FMR1*-shRNA transduced cells, we performed a subset of experiments in which 100 nM BPN-14,770 (or DMSO control) was applied for 1 day prior to recording. We found that in *Scrambled* neurons, BPN-14,770 administration unexpectedly increased AP discharge compared to vehicle. Conversely, BPN-14,770 treatment reduced AP discharge in *FMR1*-shRNA transduced neurons (Figure 3A and 3B). We found that key physiological parameters were unaffected following BPN-14,770 administration, such as: rheobase (Figure 3C), AP voltage threshold (Figure 3D) and AP maximum rise-rate (Figure 3E). However, AP maximum decay-rate displayed a strong interaction of group and treatment, indicative that differential effects of BPN between *Scrambled* and *FMR1*-shRNA neurons may be due to effects on K^+^ channel activity (Figure 3F).

**Figure 3.**
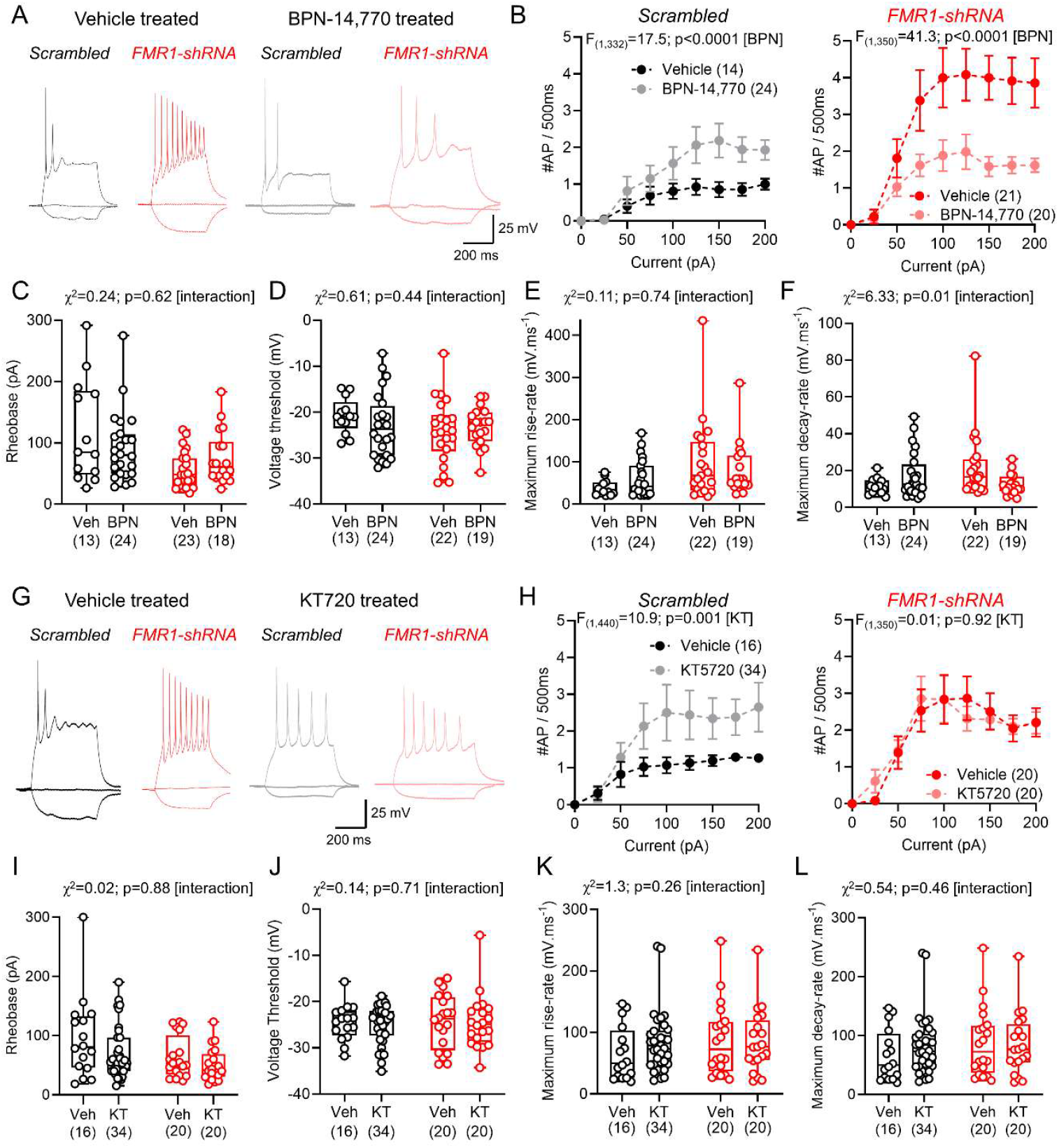
The PDE4D inhibitor BPN-14,770 corrects cortical excitability in FMRP lacking human slice cultures, while PKA increases activity in Scrambled neurons. A) example voltage responses of *Scrambled* and *FMR1-*shRNA transduced neurons following treatment with vehicle (DMSO; black and red, respectively) or the PDE4D inhibitor BPN-14,770 (100 nM; grey and pink, respectively) from 5-7 patients (Supplementary Table 3). B) AP output to depolarising current steps from *Scrambled* (left) or *FMR1*-shRNA (right) transduced neurons treated with either vehicle or BPN-14,770. Measurement of key properties of human slice culture neurons, including rheobase current (C), voltage threshold (D), maximum AP rise-rate (E), and maximum AP decay rate (F), for vehicle and BPN-14,770 treated human slice cultures. G) voltage responses of *Scrambled* and *FMR1*-shRNA transduced neurons following treatment with vehicle (DMSO; black and red, respectively) or the PKA inhibitor KT 5720 (100 nM; grey and pink, respectively) from 5 patients (Supplementary Table 4). H) AP output to depolarising current steps from *Scrambled* or *FMR1*-shRNA transduced neurons treated with either vehicle or KT 5720. Rheobase (I), voltage threshold (J), maximum AP rise-rate (K), and maximum AP decay rate (L), for vehicle and KT 5720 treated human slice cultures. Data is shown as either mean ± SEM (B, H) or box plots, depicting the median with 25-75% quartile range, and maximum & minimum. Data from individual transduced neurons shown overlaid and the number of cells shown in parenthesis. All statistical tests were performed using 2-way ANOVA (B, H) and LMM or GLMM with type-3 ANOVA, and reported above graphs.

As cAMP can modulate neuronal excitability through PKA dependent phosphorylation of sodium and potassium channels ^23,24^, we next asked whether the changes in excitability were abolished by pre-applying the PKA inhibitor KT5720 ^25^. In *Scrambled* neurons, pre-application of KT5720 led to a robust increase of AP discharge in *Scrambled* neurons selectively, but not those lacking FMRP (Figure 3G and 3H). This excitability was associated with no statistically significant interaction of group and treatment of first AP properties (Figure 3I-L). These data confirm cAMP is crucial to the function of adult human neurons. The pronounced effect on *Scrambled* neurons AP discharge may reflect high cAMP present in modified culture media ^26^, which may be occluded by impaired cAMP metabolism when FMRP is absent.

Together, these data show that neuronal hyperexcitability mediated by a loss of FMRP is likely mediated by cAMP-dependent mechanisms. This data confirms that PDE4D inhibition is a viable method to correct cortical function, and that protein-protein interactions leading to neuronal excitability deficits in the absence of FMRP are likely minimally important in human neurons.

## Discussion

In the present study we demonstrate that loss of FMRP in adult neurons leads to increased excitability, despite unaltered synaptic properties. This results from enhanced AP kinetics of neurons, which is largely corrected by administration of the PDE4D inhibitor BPN-14,770 in *FMR1*-shRNA transduced cultures. Furthermore, we show that *Scrambled* control human neurons displayed PKA-dependent inhibition of AP discharge, indicating possible refinement of this model for future study. Our data suggest a key role for FMRP in maintaining neuronal excitability in the physiological range throughout life, which are independent of the neurodevelopmental program.

### FMRP depletion leads to increased neuronal excitability in adult human neurons

In rodent models of FXS, loss of FMRP has been shown to lead to enhanced neuronal excitability of somatodendritic ^2,3,15-17,21,27^ and axonal ^7,28-30^ compartments. Indeed, such changes in excitability have been postulated to originate from compensatory mechanisms during neurodevelopment ^14^ or through ongoing protein-protein interaction which may underlie many of the endophenotypes of FXS ^8^. Determining how reduced FMRP expression alters neuronal function in adult neurons is crucial to define whether it displays a neurodevelopmental or neuromaintenance function on cellular hyperexcitability. Our data indicates that loss of FMRP in adult neurons leads to increased neuronal excitability in a cell-autonomous manner, supporting the idea that FMRP is consistently required by neurons throughout life to maintain circuit function – as has been observed for other genes associated with neurodevelopmental disorders ^31^. Specifically, we show that loss of FMRP leads to increased AP kinetics, independent of membrane resistance, a feature that has previously been attributed to altered function of the HCN-mediated currents in *FMR1*-KO mice ^2,15^. The fact that administration of BPN-14,770 returned neuronal excitability to control levels, while not reducing input resistance (an expected effect of increased cAMP activity at HCN channels) does not support these rodent studies, but indicates interactions of FMRP with ion-channels function contributing to neuronal excitability in human cortical neurons.

We found that depletion of FMRP from adult human brain circuits led to a dramatic alteration in AP kinetics, chiefly the rise and decay rates, and duration. These features are consistent with changes in functional properties of neurons we have previously observed in *FMR1*-KO mice ^7^. Furthermore, previous *in vitro* studies of primary cell-cultures from *FMR1*-KO mice reveal a shift from multi-spiking to single-spiking properties following induction of homeostatic plasticity ^32^. We find that there are more multi-spiking cells in *FMR1*-shRNA cells in humans, which indicates potential species differences in function, but how these features compensate to altered neuronal activity remains an open-ended question.

### Translational relevance of these data

Our strategy of viral depletion of FMRP using a ubiquitous promoter, aimed to mimic FXS such that *FMR1* is silenced independent of cell-type. Indeed, our transduced cells were 40-50% neurons, consistent with the general composition of the neocortex (∼42% neurons; ^33^). Nevertheless, our viral strategy achieved relatively sparse transduction rates which likely does not reflect full mutation FXS, but rather a mosaic loss of FMRP, a feature that is common in affected individuals due to the X-linked nature of *FMR1* ^34^. In this study we intentionally only examined the effects of FMRP in transduced cells. However, defining how such loss affects the function of neurons with typical levels of FMRP, or indeed wider circuit function remains unexplored. This is particularly pertinent in the context of glial expression of *FMR1*-shRNA, as selective loss of FMRP in astrocytes is known to lead to burst like activity of human neurons derived from iPSCs^35 35^. Future studies employing viral strategies targeting specific promoter sequences ^36^ may allow determination of cell-type specific effects on circuit activity.

Our data have a number of key implications for the treatment of FXS. Chiefly, that loss of FMRP in mature adult neurons leads to the emergence of hyperexcitability that has previously been attributed to neurodevelopmental cascades in rodent models ^2,3,7,16,27^. Our data shows that FMRP may serve a key neuromaintenance function, as described for other neurodevelopmental disorders (e.g. MeCP2 in Rett Syndrome ^37^). This has real -world implications for FXS treatment, as any intervention may have to be administered long-term, or gene-replacement strategies considered. This raises interesting questions regarding the FMRP-dependence of other described phenotypes, such as metabotropic glutamate receptor signalling, where sustained rescue of behavioural dysfunction was observed following lovastatin treatment ^38^. This indicates that perhaps excessive protein synthesis, as described in FXS ^39^, could be a consequence of neuronal hyperexcitability, rather than a core phenotype *per se*. Whether this is the case, and if it extends to other receptor systems known to be altered (e.g. GABA receptor signalling; ^40^) remains unknown and requires further investigation.

### Limitations of the current study

Given the relatively low number of cells transduced in human slice cultures, and their varied cell type, precise quantification of the explicit levels of FMRP in transduced cells was limited. Our estimates of FMRP depletion are likely an underestimate, as we observed high levels of non-specific autofluorescence in human slice cultures, as seen in primary antibody free controls from acute tissue (data not shown). Future studies using complementary approaches in either iPSC cultures or rodent slice cultures may validate this, in a more readily available bioresource. Second, our use of the U6 promotor for *FMR1-*shRNA expression targeted all cells, independent of type. All our recordings targeted L2/3 of the human cortex, but we cannot exclude the inclusion of diverse glutamatergic or GABAergic cells ^41,42^ which may add to the variability of our dataset. Future study using cell-type specific promoters or enhancer elements ^36^ may allow for more precise cell-type specific targeting. Finally, while our data provides the first compelling evidence for a lifelong requirement of FMRP for correct neuronal function, it is not perfect model of FXS. Affected individuals display reduced FMRP expression from early embryonic development ^34^ and while our data supports the use of BPN-14,770 in adulthood for treatment of FXS ^22^, how it affects circuits that have developed fully in the absence of FMRP and have undergone numerous compensation events during early brain development remains unexplored ^14^.

In summary, we provide evidence that FMRP depletion in adult human neurons leads to elevated neuronal excitability, which is corrected by inhibiting PDE4D, indicating the therapeutic potential of this drug in FXS.

## Methods

### Human tissue collection

Human slice cultures were generated from neocortical access tissue from patients undergoing tumour resection surgery (that would normally be disposed of during surgery), with ethical approval from the Lothian NRS Bioresource (REC number: 15/ES/0094, IRAS number: 165488) under approval number SR1319; informed consent of patients was obtained using the Lothian NRS Bioresource Consent Form. Additional approval was obtained for receiving data on patient sex, age, reason for surgery and brain region provided (NHS Lothian Caldicott Guardian Approval Number: CRD19080). Key patient demographic details are listed in Supplementary Table 1.

### Human slice cultures

Human slice cultures were prepared as previously described ^13^. Excised neocortical tissue was immediately placed in sterile ice-cold, oxygenated, and 0.22 μm filtered artificial cerebrospinal fluid (aCSF) containing (in mM): 87 NaCl, 2.5 KCl, 10 HEPES, 1.62 NaH_2_PO_4_, 25 glucose, 129.3 sucrose, 1 Na-Pyruvate, 1 Na-ascorbate, 1 thiourea, 7 MgCl_2_, and 0.5 CaCl_2_. The tissue was mounted in 2% agar, before being glued (cyanoacrylate) to a vibratome stage. 300 µm-thick slices were then cut in ice-cold and oxygenated aCSF and then sub-dissected into slices containing cortical columns.

Slices were placed into 0.22 μm-filtered wash buffer composed of oxygenated Hanks Balanced Salt Solution (HBSS, Thermo Fisher: 14025092), HEPES (20 mM) and 1x penicillin/streptomycin (Thermo Fisher: 15140122) with osmolarity of 305 mOsm, pH 7.3 for 20 min at room temperature. Slices were then plated on membranes (Millipore: PICM0RG50) sitting on top of 750 μL of a second wash medium, placed inside 35 mm culture dishes. The second wash medium was 0.22 µm-filtered and composed of 96% BrainPhys^®^ Neuronal Medium (StemCell Technologies: 5790) supplemented with 1x N2 (Thermo Fisher: 17502001), 1x B27 (Thermo Fisher: 17504044), 40 ng/mL hBDNF (StemCell Technologies: 78005), 30 ng/mL hGDNF (StemCell Technologies: 78058), 30 ng/mL Wnt7a (Abcam: ab116171), 2 μM ascorbic acid, 1 mM dibutyryl cAMP (APExBIO: B9001), 1 μg/mL laminin (APExBIO: A1023), 1x penicillin/streptomycin (Thermo Fisher: 15140122), 3 U/mL nystatin (Merck: N1638) and 20 mM HEPES. Slice cultures were kept in the second wash medium in an incubator at 37 °C with 5% CO_2_ for at least 1 hour, after which the medium was aspirated and replaced with maintenance medium (as second wash medium, but with no HEPES). 100% medium exchanges occurred on DIV 3 and 7, until collection at days DIV 8-10.

### Viral transduction of FMR1 shRNA and BPN-14,770/KT5720 treatment

Viral transduction was carried out using two viral vectors-Scrambled control and shRNA-*Fmr1*. Each construct was packaged into a lentiviral vector expressing both the U6 promotor and enhanced green fluorescent protein (eGFP). Final titres of each virus were >10x^8^ TU/ml. Both viruses were purchased form VectorBuilder.com (Scrambled: VB010000-0009mxc; shRNA-Fmr1: VB900043-0662ekc). Treatment of the cultures consisted of adding 1 µL of either scrambled or shRNA-Fmr1 atop of each slice at DIV 0. For PDE4D inhibition experiments, 100 nM BPN-14,770 or 100 nM of DMSO was added to the media for 1 day prior to recording. For KT5720 treatment, slice cultures were dosed with 100 nM of KT5720 or 100 nM DMSO for 2-4 hours before recording. During recording, slices were continuously perfused with carbogenated ACSF with 100 nM KT5720/DMSO.

### Electrophysiological recording

For electrophysiological recording, slice cultures were transferred to a submerged recording chamber perfused with carbogenated ACSF (in mM: 125 NaCl, 2.5 KCl, 25 NaHCO_3_, 1.25 NaH_2_PO_4_, 25 glucose, 1 MgCl_2_, 2 CaCl_2_) maintained at near physiological temperatures (31 ± 1 °C) with an inline heater (LinLab, Scientifica, UK) at a flow rate of 6–8 ml/min. Slices were visualized with IR-DIC illumination (Slicescope, Scientifica, UK) using a 40x objective lens (N.A. 1.0, Fluoplan, Olympus). Whole-cell patch-clamp recordings were made with a Multiclamp 700B amplifier (Molecular Devices, USA). Recording pipettes were pulled from borosilicate glass capillaries (1.5 mm outer/0.86 mm inner diameter, Harvard Apparatus, UK) on a horizontal electrode puller (P-1000, Sutter Instruments, CA, USA), which when filled with intracellular solution gave a pipette resistance of 5-7 MΩ. All signals were filtered online at 10 kHz using the built in 4-pole Bessel filter of the amplifier, digitized at 20 kHz on an analogue-digital interface (Digidata 1440, Axon Instruments, CA, USA), and acquired with pClamp software (pClamp 10, Axon Instruments, CA, USA). Data were analysed offline using the open source Stimfit software package (http://www.stimfit.org). Cells were rejected if the membrane potential was more depolarised than −40 mV, series resistance >30 MΩ, input resistance >1000 MΩ, or the series resistance changed by more than 20% over the course of the recording.

To characterise intrinsic physiology of neurons, we first applied a -10 pA current step (500 ms duration, 30 repetitions) from resting membrane potential – to allow measurement of input resistance and membrane time-constant. To test AP discharge properties and excitability, we applied a family of 500 ms hyper-to depolarising current steps using either -100 to +400 pA (25 pA steps) or -25 to +200 pA (5 pA steps), depending on cell resistance. For membrane impedance and resonant frequency, we delivered a sinusoidal “chirp” of 0.1 – 20 Hz frequency of either 50 pA or 100 pA (20 second duration). All AP parameters were measured from the first AP elicited above rheobase from -70 mV, maintained by applying a bias current. AP kinetics were measured from threshold (10 mV/ms). For analysis, the number of APs was measured per current step at a threshold of -15 mV and AP kinetics calculated from the first response that crossed 0 mV. For chirp stimuli, we first ran a fast-Fourier transform of the traces, then measured the peak response, giving the resonant frequency and impedance). To measure synaptic inputs, we recorded 2 minutes of activity in voltage-clamp from a holding potential of -70 mV and Spontaneous excitatory post-synaptic currents (sEPSC) were detected using a template-matching algorithm, with events included if they were 3x larger than the standard deviation of preceding (5 ms) event-free baseline.

### Immunohistochemisty

To confirm knock-down of FMRP and identify transduction efficiency in different cell populations, we performed immunohistochemical analysis of fixed human slice cultures. For this, slices were fixed at DIV7-10 overnight in 4% paraformaldehyde in 0.1 M phosphate buffer (PB), then transferred to PBS. For labelling, human slice cultures were washed in PBS, blocked for 1 hr in PBS with 10% normal goat serum and 0.3% Triton X-100 and then incubated overnight at 4 °C with antibodies recognising MAP2 (1:500, guinea pig Synaptic Systems: 188004) and FMRP (rabbit 1:1000, Abcam, ab17722, LOT: 1080719-1), in a solution containing 5 % normal goat serum and 0.3 % Triton X-100, and 0.05 % NaN_3_ diluted in PBS. The slices were then washed 3 × in PBS, before incubation for 2 h in 1:500 secondary antibodies, namely: goat anti-guinea pig 568 (2 μg/ml; Thermo Fisher), goat anti-rabbit 568 (2 μg/ml; Thermo Fisher), goat anti-rabbit 647 (2 μg/ml; Thermo Fisher) and streptavidin conjugated to AlexaFluor 633 nm (2 μg/ml; Thermo Fisher) which were diluted in PBS containing 3% NGS, 0.1% Triton-X100, and 0.05% NaN_3_ and incubated overnight at 4 °C. Slices were washed 3 × in PBS, then 2× in PB and then mounted on slides in antifade mounting medium (either VectaShield, H-1900 or Fluoromount, Southern Biotech). Images were taken using 20x (1.4 N.A.) or 63x (1.4 N.A.) objectives on a Leica SP8 confocal microscope. Images were acquired at either 1024 x 1024 μm (20x) or 2048 x 2048 μm (63x) pixel resolution at 0.30 or 0.13 μm z-steps. respectively. All images were analysed offline using FIJI/ImageJ.

### Statistical analysis

All data are presented as the mean ± SEM with data from individual cells shown overlaid. To account for inter-individual variability, data were analysed with a linear mixed-effects model (LMM) or generalised form (GLMM) in R software using:

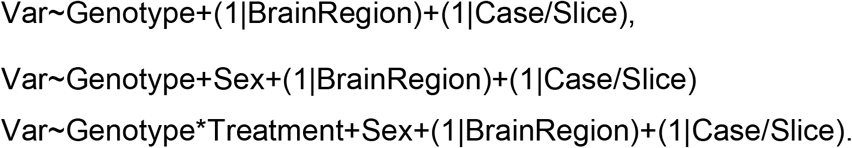

Probability distributions for models were chosen by goodness of fit to normal, log-normal or gamma distributions. Viral treatment, drug treatment, sex and interactions were used as fixed effects, while case, slice and brain region were selected as random effects. When within case-datasets are shown (Figure 1), they were tested for normality (d’Agostino-Pearson test) and Mann–Whitney non-parametric U-tests or Wilcoxon signed-rank tests performed. Statistically significant differences were defined as p < 0.05. Statistical tests employed are indicated throughout the text. GraphPad Prism or R studio were used for all statistical analyses.

## Supporting information

Supplementary Materialas

## Acknowledgements

We thank Ania Sumera and Jamie Elliott for help with data analysis approaches and proof-reading reading the draft manuscript. This work would not be possible without the support of the EMERGE research nurse team and the patients who provided consent for their brain tissue to be used for research. This work is funded by UK Medical Research Council (MR/Y014529/1-SAB), Simons Initiative for the Developing Brain (SAB), Race Against Dementia (ARUK-RADF-2019a-00 – CSD), and the Dyson Foundation (CSD)

