## Supplementary Materialas for "FMRP regulates adult human cortical neuron excitability via cyclic-AMP signalling"

Supplementary Materials

| **Sex** | **Age** | **Brain region** | **Seizure history?** |
|---|---|---|---|
| M | 20-25 | Temporal | Yes |
| F | 25-30 | Frontal | No |
| F | 35-40 | Frontal | No |
| M | 40-45 | Frontal | No |
| M | 45-50 | Parietal | Yes |
| M | 50-55 | Temporal | No |
| M | 55-60 | Temporal | Yes |
| M | 55-60 | Parietal | No |
| M | 55-60 | Temporal | No |
| M | 60-65 | Occipital | No |
| M | 60-65 | Temporal | No |
| M | 60-65 | Temporal | No |
| M | 60-65 | Temporal | No |
| M | 65-70 | Temporal | No |
| M | 65-70 | Temporal | No |
| F | 65-70 | Parietal | Yes |
| M | 65-70 | Occipital | Yes |
| M | 65-70 | Frontal | No |
| M | 75-80 | Temporal | Yes |
| M | 75-80 | Frontal | Yes |
| M | 65-70 | Frontal | No |
| M | 65-70 | Parietal | No |
| F | 55-60 | Temporal | Yes |
| M | 20-25 | Frontal | Yes |
| M | 60-65 | Parietal | No |

*Supplementary table 1: Key clinical features of patients from whom tissue was cultured.* A list of key patient information, with age range, biological sex, and seizure history indicated.

| Electrophysiological Property | *Scrambled* 23 cells/8 cases | *FMR1*-ShRNA 33 cells/12 cases | $\chi^2$ ; P |
|---|---|---|---|
| Membrane potential (mV) | -62.4 ± 2.9 | -60.5 ± 1.9 | 0.465 ; 0.495 |
| Input resistance (MΩ) | 415 ± 46 | 460 ± 49 | 1.74 ; 0.187 |
| Membrane time-constant (ms) | 50.4 ± 6.1 | 52.1 ± 6.3 | 0.081 ; 0.776 |
| Capacitance (pF) | 146 ± 20 | 141 ± 21 | 0.46 ; 0.500 |
| Rheobase (pA) | 104 ± 15 | 60 ± 8 | 10.9 ; 0.0009*** |
| Voltage threshold (mV) | -23.6 ± 1.4 | -25.5 ± 0.9 | 2.24 ; 0.135 |
| AP amplitude (mV) | 86.7 ± 3.4 | 95.9 ± 2.5 | 5.25 ; 0.022* |
| AP 20-80% rise-time (ms) | 0.7 ± 0.1 | 0.5 ± 0.05 | 5.17 ; 0.023* |
| AP half-height width (ms) | 3.8 ± 0.3 | 2.9 ± 0.2 | 6.50 ; 0.01* |
| AP maximum rise ($mV.ms^{-1}$) | 68 ± 13 | 116 ± 10 | 10.6 ; 0.001** |
| AP maximum decay ($mV.ms^{-1}$) | 10 ± 1 | 21 ± 3.0 | 12.7 ; 0.0003*** |
| FI Slope ($AP.\ pA^{-1}$) | 0.04 ± 0.01 | 0.1 ± 0.03 | 9.55 ; 0.002** |
| Peak firing (AP/500ms) | 2 ± 0.3 | 5 ± 1 | 5.42 ; 0.01* |

*Supplementary Table 2: Electrophysiological properties of FMR1-shRNA cells compared with Scrambled controls.* Key intrinsic physiological measurements made from human slice cultures, including AP kinetics. Data is shown as mean ± SEM. Statistics shown as p-values from linear mixed effects models (LMM; Var~Genotype+Sex+(1|Brain Region)+(1|Case/Slice)) comparing the effect of genotype. Number of cells and cultures contributing to each dataset are indicated.

| Electrophysiological Property | DMSO vehicle controls: *Scrambled* 13 cells 5 cases | DMSO vehicle controls: *FMR1*-shRNA 22 cells 7 cases | 100 nM BPN-14,770: *Scrambled* 23 cells 6 cases | 100 nM BPN-14,770: *FMR1*-ShRNA 19 cells 7 cases | $\chi^2$; *p* (group) | $\chi^2$; *p* (treatment) | $\chi^2$; *p* (interaction) |
|---|---|---|---|---|---|---|---|
| Membrane potential (mV) | -60 ± 3 | -58 ± 2 | -64 ± 3 | -58 ± 2 | 4.51 ; 0.03* | 0.211 ; 0.65 | 0.610 ; 0.43 |
| Input resistance (MΩ) | 394 ± 69 | 476 ± 39 | 404 ± 41 | 544 ± 59 | 4.09 ; 0.04* | 0.975 ; 0.323 | 0.294 ; 0.59 |
| Membrane time-constant (ms) | 36 ± 4 | 51 ± 5 | 43 ± 5 | 52 ± 7 | 1.22 ; 0.27 | 0.046 ; 0.83 | 0.079 ; 0.79 |
| Capacitance (pF) | 128 ± 29 | 129 ± 22 | 116 ± 12 | 131 ± 26 | 0.648 ; 0.42 | 0.686 ; 0.41 | 0.617 ; 0.43 |
| Rheobase (pA) | 120.0 ± 23 | 55 ± 6 | 92 ± 11 | 76 ± 10 | 2.37 ; 0.12 | 1.28 ; 0.26 | 0.239 ; 0.62 |
| Voltage threshold (mV) | -21 ± 1 | -24 ± 1 | -23 ± 1 | -23 ± 1 | 0.106 ; 0.74 | 0.075 ; 0.78 | 0.606 ; 0.44 |
| AP amplitude (mV) | 82 ± 3 | 91 ± 3 | 84 ± 3 | 91 ± 3 | 4.90 ; 0.02* | 0.047 ; 0.83 | 0.046 ; 0.83 |
| AP 20-80% rise-time (ms) | 0.6 ± 0.03 | 0.5 ± 0.04 | 0.5 ± 0.04 | 0.5 ± 0.03 | 0.027 ; 0.87 | 0.283 ; 0.59 | 1.33 ; 0.25 |
| AP half-height width (ms) | 3.1 ± 0.3 | 2.3 ± 0.2 | 2.5 ± 0.2 | 3.9 ± 0.4 | 13.5 ; 0.0002*** | 17.4 ; 0.00003**** | 17.6 ; 0.00003**** |
| AP maximum rise ($mV.ms^{-1}$) | 43 ± 7 | 96 ± 20.0 | 68 ± 12 | 78 ± 14 | 1.67 ; 0.20 | 0.038 ; 0.85 | 0.106 ; 0.74 |
| AP maximum decay ($mV.ms^{-1}$) | 11 ± 1 | 22 ± 4 | 17 ± 3 | 12 ± 1 | 0.758 ; 0.38 | 7.34 ; 0.006** | 6.33 ; 0.01* |
| FI Slope ($AP. pA^{-1}$) | 0.03 ± 0.01 | 0.08 ± 0.01 | 0.05 ± 0.01 | 0.06 ± 0.01 | 0.013 ; 0.91 | 3.81 ; 0.051 | 1.27 ; 0.87 |
| Peak firing | 1 ± 0.2 | 5 ± 0.9 | 2 ± 0.3 | 2 ± 0.2 | 1.20 ; 0.19 | 13.2 ; 0.0003*** | 13.9 ; 0.0002*** |

*Supplementary Table 3: Electrophysiological properties of HOSCs treated with BPN-14,770 compared with vehicle controls.* Key intrinsic physiological measurements made from human slice cultures, including AP kinetics. Data is shown as mean ± SEM. Statistics shown as p-values from linear mixed effects models (LMM; Var~Genotype*Treatment+Sex+(1|Brain Region)+(1|Case/Slice)) comparing the effect of treatment. Number of cells and cultures contributing to each dataset are indicated.

| Electrophysiological Property | DMSO vehicle controls | | 100 nM KT5720 | | | | |
|---|---|---|---|---|---|---|---|
| | *Scrambled* 16 cells 6 cases | *FMR1*-ShRNA 20 cells 6 cases | *Scrambled* 34 cells 7 cases | *FMR1*-ShRNA 20 cells 6 cases | $\chi^2$; P (group) | $\chi^2$; P (Treatment) | $\chi^2$; P (interaction) |
| Membrane potential (mV) | -63 ± 3 | -60 ± 2 | -63 ± 2 | -60 ± 2 | 1.66 ; 0.20 | 0.102 ; 0.75 | 0.097 ; 0.76 |
| Input resistance (MΩ) | 475 ± 63 | 545 ± 62 | 507 ± 49 | 639 ± 54 | 4.22 ; 0.04* | 2.31 ; 0.13 | 0.60 ; 0.44 |
| Membrane time-constant (ms) | 46 ± 8 | 61 ± 7 | 55 ± 7 | 70 ± 9 | 5.07 ; 0.02* | 1.16 ; 0.28 | 0.028 ; 0.87 |
| Capacitance (pF) | 124 ± 27 | 134 ± 16 | 126 ± 13 | 125 ± 18 | 0.0001 ; 0.99 | 0.170 ; 0.68 | 0.160 ; 0.69 |
| Rheobase (pA) | 96 ± 17 | 64 ± 8 | 69 ± 8 | 51 ± 6 | 2.00 ; 0.16 | 1.46 ; 0.23 | 0.024 ; 0.88 |
| Voltage threshold (mV) | -25 ± 1 | -24 ± 1 | -25 ± 1 | -24 ± 1 | 0.298 ; 0.59 | 0.022 ; 0.88 | 0.135 ; 0.71 |
| AP amplitude (mV) | 86 ± 4 | 92 ± 2 | 92 ± 2 | 96 ± 3 | 1.54 ; 0.21 | 1.12 ; 0.29 | 0.12 ; 0.73 |
| AP 20-80% rise-time (ms) | 0.5 ± 0.04 | 0.5 ± 0.03 | 0.5 ± 0.02 | 0.5 ± 0.05 | 0.840 ; 0.36 | 0.836 ; 0.36 | 1.78 ; 0.18 |
| AP half-height width (ms) | 3.2 ± 0.4 | 3.7 ± 0.4 | 3.3 ± 0.3 | 3.9 ± 0.4 | 2.82 ; 0.09 | 0.294 ; 0.59 | 0.285 ; 0.59 |
| AP maximum rise ($mV.ms^{-1}$) | 66 ± 11 | 87 ± 13 | 82 ± 9 | 85 ± 12 | 0.001 ; 0.97 | 0.015 ; 0.90 | 1.27 ; 0.26 |
| AP maximum decay ($mV.ms^{-1}$) | 12 ± 1 | 14 ± 2 | 18 ± 3 | 13 ± 2 | 2.20 ; 0.14 | 0.054 ; 0.82 | 0.54 ; 0.46 |
| FI Slope ($AP. pA^{-1}$) | 0.04 ± 0.01 | 0.06 ± 0.07 | 0.06 ± 0.008 | 0.07 ± 0.01 | 0.250 ; 0.617 | 0.215 ; 0.64 | 1.35 ; 0.24 |
| Peak firing | 2 ± 0.3 | 3 ± 0.6 | 4 ± 0.8 | 3 ± 06 | 0.092 ; 0.76 | 0.223 ; 0.64 | 0.557 ; 0.46 |

*Supplementary Table 4: Electrophysiological properties of* HOSCs *treated with KT-5720 compared with vehicle controls.* Key intrinsic physiological measurements made from human slice cultures, including AP kinetics. Data is shown as mean ± SEM. Statistics shown as p-values from linear mixed effects models (LMM; Var~Genotype*Treatment+Sex+(1|Brain Region)+(1|Case/Slice)) comparing the effect of treatment. Number of cells and cultures contributing to each dataset are indicated.

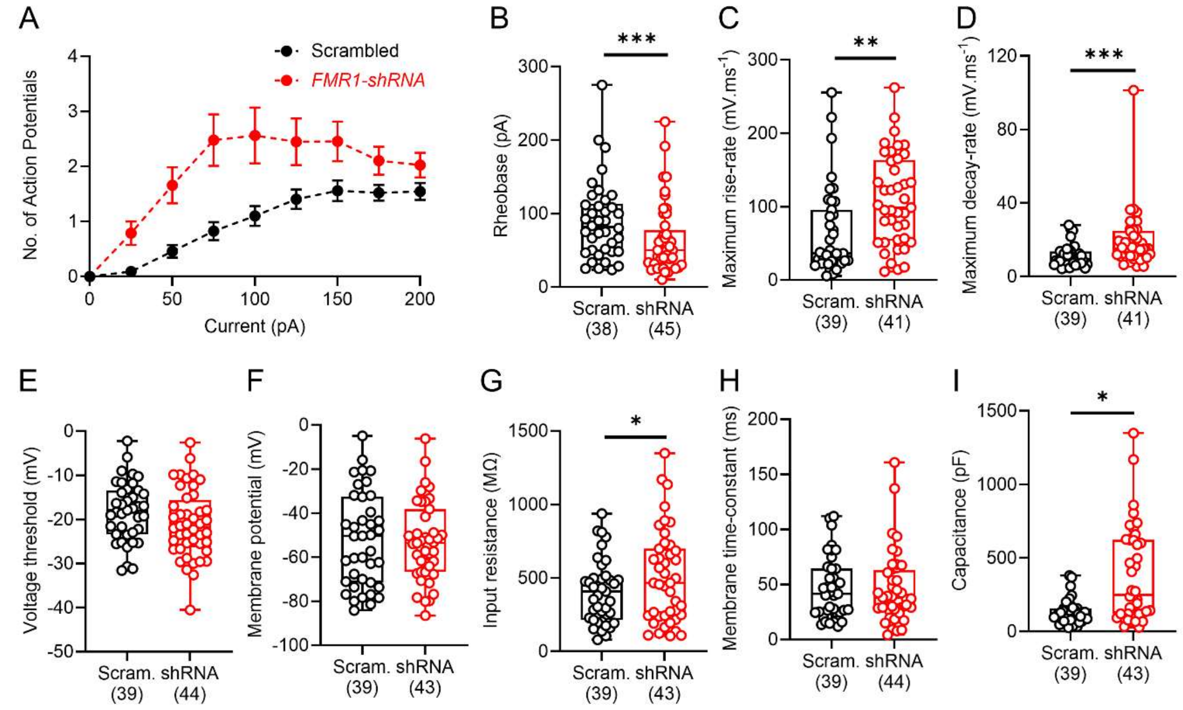


*Supplementary Figure 1: Comparison of electrophysiological parameters of all neurons recorded from human slice cultures, regardless of resting membrane potential*. *A*) AP output to depolarising current steps from human slice culture neurons recorded with any resting membrane potential. Measurement of active properties of all human slice culture neurons from *Scrambled* (black) or *FMR1*-shRNA (red) transduced neurons, including rheobase current (B), maximum AP rise-rate (C), and maximum AP decay rate (D), voltage threshold (E). Quantification of passive membrane properties of recorded neurons, including resting membrane potential (F), input resistance (G), membrane time-constant (H), and calculated capacitance (I). Data is shown as either mean ± SEM (B) or box plots, depicting the median with 25-75% quartile range, and maximum & minimum. Data from individual transduced neurons are shown overlaid. All statistical tests were performed using a LMM or GLMM with type-3 ANOVA and shown as: * – $p$<0.05, ** – $p$>0.01, *** – $p$<0.001, **** – $p$<0.0001.

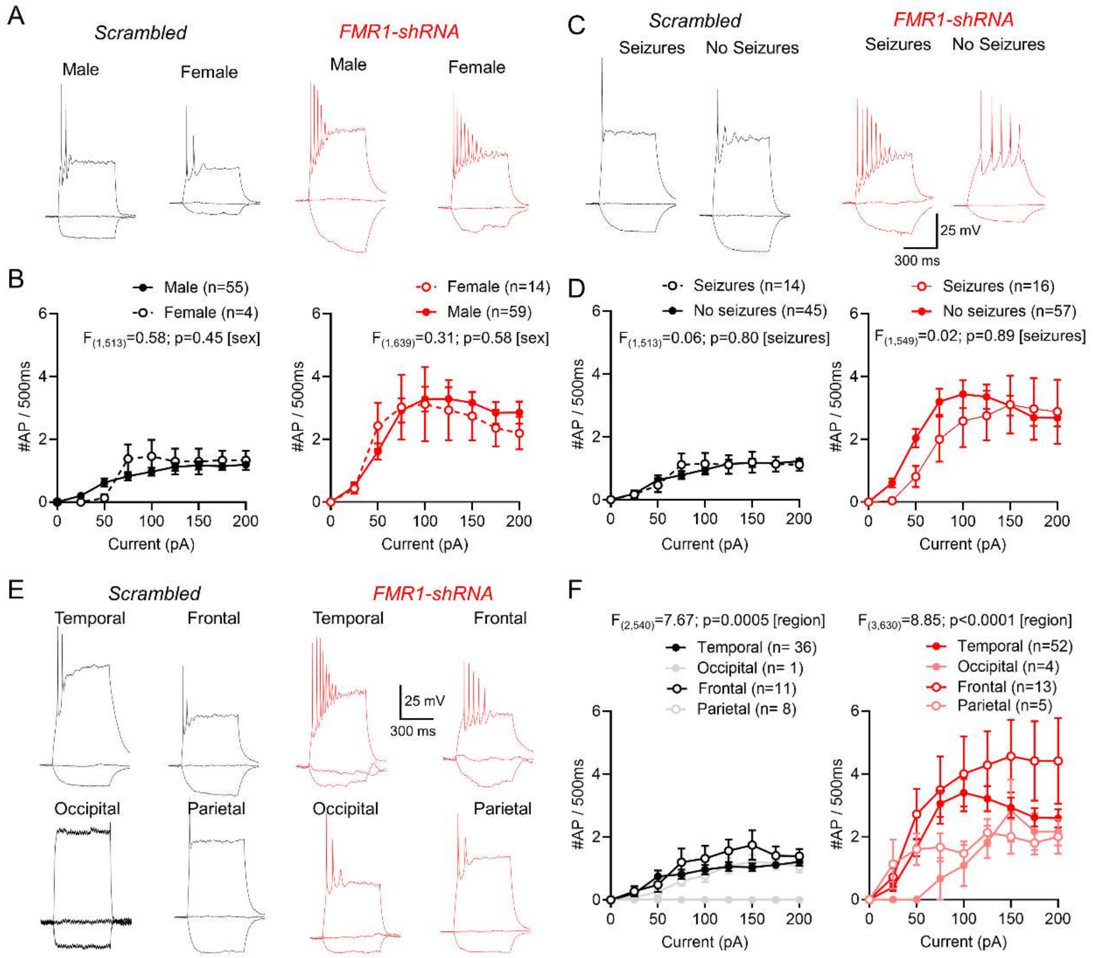


*Supplementary Figure 2: Comparison of FMR1-shRNA transduction on neuronal output, dependent on biological sex, seizure presentation, and brain region.* A) Example voltage responses of neurons in human slice cultures from *Scrambled* (black) or *FMR1*-shRNA (red) transduced neurons in response to -25, 0, and + 100 pA current steps (500 ms) from male and female patients. B) AP output to depolarising current steps from human slice culture neurons from male (filled circles) and female (open circles) from *Scrambled* (black, left) or *FMR1*-shRNA (red, right); recorded with a resting membrane potential below -40 mV. C) Voltage responses of neurons in human slice cultures from *Scrambled* or *FMR1*-shRNA transduced neurons under the same conditions as (A), but from patients who had experienced seizures or not. D) AP output to depolarising current steps from human slice culture neurons from patients without seizures (filled circles) or those who experienced seizures (open circles) from *Scrambled* or *FMR1*-shRNA neurons. E) Voltage responses of neurons in human slice cultures from *Scrambled* or *FMR1*-shRNA transduced neurons under the same conditions as (A), but from tissue resected from temporal (filled black/red circles), occipital (filled grey/pink circles), frontal (open black/red circles), or parietal (open grey circles) cortices from Scrambled and FMR1-shRNA patients. Note the variability of excitability of neurons dependent on brain area sampled. Data is shown as mean ± SEM with data from individual transduced neurons shown overlaid. All statistical tests were performed using a 2-way ANOVA, and reported above graphs.

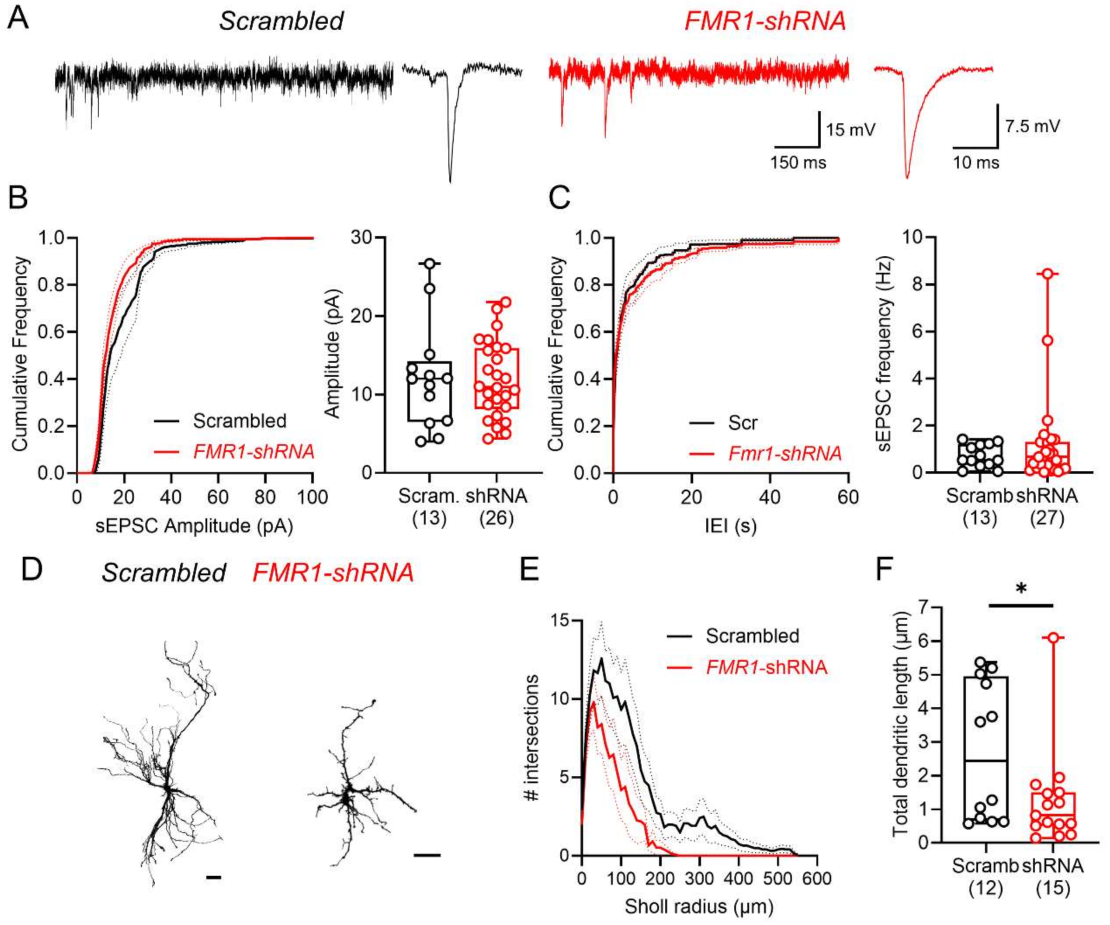


*Supplementary Figure 3: Synaptic properties of human slice culture neurons are unaffected following loss of FMRP, but neurons display morphological impairments.* A) Example voltage-clamp recordings from -70 mV from transduced neurons in Scrambled (black) and *FMR1*-shRNA (red) human slice cultures. Inset, average spontaneous EPSC (sEPSC) responses from the same recordings. B) quantification of sEPSC amplitudes shown as average cumulative distributions (left) and box-plots (right) from all recorded cells ($\chi^2$=0.08, p=0.77, LMM). C) sEPSC inter-event intervals (IEI) and average frequency ($\chi^2$=0.13, p=0.72, LMM), plotted in the same manner. D)

Example reconstructions of biocytin-filled transduced neurons from Scrambled and *FMR1*-shRNA. Scale bars: 100 µm. E) Sholl analysis of the reconstructed cells, revealing reduced dendritic complexity in FMR1-shRNA transduced neurons. F) quantification of total dendritic length in all reconstructed neurons revealed shorter dendritic arborisations following loss of FMRP ($\chi^2$=5.02, p=0.024, LMM). Data is shown as either average cumulative distributions (B, C) mean ± SEM (E) or box plots, depicting the median with 25-75% quartile range, and maximum & minimum (B, C, F). Data from individual neurons are shown overlaid. All statistical tests were performed using a LMM or GLMM with type-3 ANOVA and shown as: * – $p<0.05$.
